# Antibiotic-resistant *Salmonella species* and *Escherichia coli* in broiler chickens from farms, abattoirs and open markets in selected districts of Zambia

**DOI:** 10.1101/2020.04.20.050914

**Authors:** Nelson Phiri, Geoffrey Mainda, Mercy Mukuma, Ntazana N. Sinyangwe, Luke John Banda, Geoffrey Kwenda, Elizabeth Muligisa-Muonga, Bumbangi Nsoni Flavien, Mwaba Mwansa, Kaunda Yamba, Musso Munyeme, John Bwalya Muma

## Abstract

*Salmonella* species and *Escherichia coli* are major bacterial enteropathogens of global public health importance that cause foodborne diseases, thereby contributing to increased human morbidity and mortality. Both pathogens have also been found to contribute towards the spread of antimicrobial resistance through the food chain, especially in poultry. The aim of this study was to determine the occurrence of antibiotic-resistant *Salmonella sp*. and *E. coli* in broiler chickens at farm level, abattoirs and open markets in selected districts of Zambia. A cross-sectional study was undertaken in seven districts of Zambia to determine the resistance profiles of *Salmonella sp*. and *E. coli* obtained from broiler chickens at farms, abattoirs and open markets. A total of 470 samples were collected, including litter, cloacal swabs and carcass swabs. Samples were inoculated into buffered peptone water, sub-cultured onto MacConkey and Xylose Lysine Deoxycholate agar plates. Identification of *Salmonella sp. and E. coli* was done using the API-20E kit and confirmation by 16S rDNA sequencing. Confirmed isolates were tested against a panel of 10 antibiotics using the Kirby-Bauer disc-diffusion method and interpreted according to the Clinical Laboratory Standards Institute guidelines. Analysis of the antibiotic susceptibility test results was done using WHONET 2018 software. Overall, 4 *Salmonella spp*. and 280 *E. coli* were isolated. One of the *Salmonella sp*. was resistant to ampicillin (25%), amoxicillin/clavulanic acid (25%) and cefotaxime (25%). *E. coli* antibiotic resistance was highest to tetracycline (81.4%) and lowest to imipenem (0.7%). The antibiotic susceptibility profile revealed 55% (154/280) multidrug resistant *E. coli*, with the highest multidrug resistance profile (20.7%) in the ampicillin-tetracycline-trimethoprim/sulfamethoxazole drug combination. Furthermore, 4.3% (12/280) of the isolates showed Extensive Drug resistance. The levels of antimicrobial resistance to *E. coli* and *Salmonella* observed in market-ready chickens is of public health concern.

## Introduction

Poultry production is one of the most important activities in the livestock sector in many countries, including Zambia. Production of chicken meat requires great care to assure food safety. Disease burden has however, remained a great challenge in poultry production (1). Some of the common microbial pathogens isolated from fresh poultry meat include *Salmonella sp*., *Campylobacter spp*. and *Escherichia coli* (2). Failure to manage these pathogens in poultry has led to various food-borne disease outbreaks in countries such as South Africa and Botswana (3, 4), as well as in the United States (5).

Even though progress is being made in the control of these pathogens, they tend to evolve and generate new challenges such as antibiotic resistance (6). Antibiotic usage is considered as one of the most important factors in promoting the emergence, selection and dissemination of antibiotic-resistant microorganisms in both veterinary and human medicine (7). Antibiotic usage selects for resistance in pathogenic bacteria and the endogenous bacterial flora of exposed animals and humans (8). Also, resource-constrained countries face challenges that co-exist and facilitate the spread of bacteria during livestock production, transportation and processing. These challenges include high bird population density in poultry houses and/or poor infection control measures such as lack of vaccinations and biosecurity (9).

In Zambia, processing plants such as abattoirs are still facing challenges in producing wholesome and safe food of animal origin for human consumption due to contaminations by antibiotic-resistant bacteria (10). This is partly because some poultry farmers are using antibiotics as growth promoters, which are perceived as an inexpensive management practice (11), while other farmers use antibiotics in disease prevention as a mitigation measure against the highly prevalent unhygienic conditions and absence of biosecurity (12). Consequently, antibiotics are found in meat as residues and bacteria are continuously being exposed to them with a risk of developing resistance (13). This study was, therefore, carried out to determine the occurrence of antibiotic-resistant *Salmonella sp*. and *E. coli* isolated from broiler chickens that are intended for human consumption at farm level, abattoirs and open markets in selected districts of Zambia.

## Materials and methods

### Study design, site and population

A cross-sectional study was conducted from December 2017 to June 2018 to investigate the occurrence of antibiotic-resistant *Salmonella sp*. and *E. coli* in broiler chickens from poultry farms, commercial abattoirs and open markets. Litter and cloacal swab samples were collected from 7 districts: Chilanga, Chongwe, Kafue, Lusaka (Lusaka Province), Choma (Southern Province), Kabwe (Central Province) and Kitwe (Copperbelt Province). In Lusaka Province, only two commercial poultry abattoirs gave consent to the study. In Choma, Kitwe and Kabwe, where no poultry abattoirs were available, freshly voided faecal droppings from market-ready broiler chickens and cloacal swab samples were collected from farms and open markets. Chickens that were condemned at slaughter or point of sale were excluded from the study.

### Sample size and sampling technique

#### Poultry houses

In all the districts included in the study, there was no information on the number of farmers who reared broiler chickens as most of whom were seasonal farmers. Seasonal farmer was defined as the farmer who keep broiler chicken when the production parameters including cost of feed, cost of medicines are favourable and stops when they are not. Therefore, the snowball sampling method was used, and farmers in production were initially identified with the help of a local veterinary assistant or livestock officer. Such farmers would then lead to other farmers in season of production. At each farm, several poultry litter portions (one sample per 25m^2^) were collected from each poultry house and pooled for laboratory analysis. Using this technique, a total of 212 pooled litter samples were collected from the following districts: Chilanga (n=31), Chongwe (n=23), Kafue (n=33), Lusaka (n=24), Choma (n=17), Kabwe (n=39) and Kitwe (n=45).

#### Abattoirs

A total of two abattoirs were included in this study. Three cloacal and three carcass swabs were collected from each batch of chickens supplied to each of the abattoirs since only 25 farmers supplied chickens during the period of study. Ten (10) and fifteen (15) chickens and cloacal swabs were sampled from abattoir A and B, respectively. The two (2) were the main abattoirs in the study area and supplied poultry meat to supermarkets and open markets throughout the country. Random “blind” sampling method was used to select the 3 chickens and cloacal swabs. This method was used as it yields information about the average composition of the lot. It is employed when there is no information or method for determining which units (bacterial pathogens) are violated (14). A total number of 150 samples were collected from the two abattoirs, comprising 75 cloacal swabs collected in the receiving bay before hosting the birds on the hackles (targeting bacteria originating from farms) and 75 carcass swabs collected during the packaging process before the carcasses were chilled (to ascertain the efficiency of processing and cross-contamination). Carcass swabs were collected from under the wings of the chicken where the bacterial population is thought to concentrate during processing (15).

#### Open Markets

Choma, Kabwe and Kitwe districts did not have any abattoir at the time of sampling. Therefore, only broiler chickens sold on open markets were available for cloacal swab collection. Samples were collected from chickens of all vendors available on the day of the visit. Random “blind” sampling method was equally used at these sites. A total of 108 cloacal swabs were collected with the following distribution: Choma (35), Kabwe (40), and Kitwe (33).

All samples were immediately transferred into Amie’s transport media (Oxoid, Basingstoke, UK) in a cool box with ice packs and transported to the Public Health Laboratory at the University of Zambia, School of Veterinary Medicine for analysis. Samples were processed and analyzed within 24 hours of collection

### Laboratory analysis

Laboratory analysis included isolation of *Salmonella sp*. and *E. coli*, identification, confirmation of the isolates and antibiotic susceptibility testing (AST). Laboratory protocols for bacterial isolation recommended by the Food and Drug Administration’s Bacteriological Analytical Manual were used with few modifications (14, 16). All media used were prepared according to manufacturer’s instructions. The media were quality controlled using control strains *E. coli* ATCC 25922 and *Salmonella typhimurium* ATCC 14028.

### Isolation and identification of *Salmonella species*

Litter and swabs samples were pre-enriched in 10 mL buffered peptone water (Oxoid, Basingstoke, UK) and incubated at 37°C for 24 hours. Aliquots from the pre-enrichment broth were inoculated into Rappaport Vassiliadis medium (Oxoid, Basingstoke, UK), a selective enrichment medium for *Salmonella sp*., at a ratio of 1:10 and incubated at 37°C for 48 hours. A loop full of enriched broth was streaked on Xylose Lysine Deoxycholate (XLD) agar plates (Oxoid, Basingstoke, UK) and incubated aerobically at 37 °C for 18-24 hours. The presumptive identification of *Salmonella sp*. was based on morphological characteristics of colonies of non-lactose fermenters. Suspected colonies of *Salmonella sp*. from each plate were subjected to serological testing using polyvalent serum against O and H antigens. Presumptive *Salmonella sp*. colonies were then sub-cultured on nutrient agar plates (Oxoid, Basingstoke, UK), incubated at 37°C for 18 to 24 hours, and the resulting pure colonies subjected to biochemical identification using the API-20E test kit (bioMérieux, Marcy I’Etoile, France) according to the manufacturer’s instructions. The identity of the isolates was confirmed by sequencing of the bacterial 16S rDNA molecule (17).

### Isolation and identification of *E. coli*

For the isolation of *E. coli*, litter and swabs samples were placed in 10mL of buffered peptone water (Oxoid, Basingstoke, UK) as a pre-enrichment media and incubated at 37°C for 24 hours. Aliquots from the pre-enrichment broth were sub-cultured onto MacConkey agar plates (Oxoid, Basingstoke, UK) and incubated aerobically for an additional 18-24 hours at 37°C. Lactose fermenting colonies were then sub-cultured onto Eosin Methylene Blue (EMB) agar plates (Oxoid, Basingstoke, UK) and incubated aerobically at 37°C for 18-24 hours. After incubation, presumptive *E. coli* colonies were observed to have a distinct green metallic sheen and confirmed by using the API-20E test kit and 16S rDNA sequencing as described for *Salmonella* isolates. All isolates were placed in 10% glycerol and stored at −20°C for a short period until AST was done.

### Antibiotic sensitivity testing

The antibiotic susceptibility testing was done using the Kirby-Bauer disc diffusion method on Müeller-Hinton agar plates (Oxoid, Basingstoke, UK) (18). Cell suspension densities equal to 0.5 McFarland turbidity were prepared from fresh, pure cultures of either *Salmonella sp*. or *E. coli* isolates grown overnight using a Nephelometer. Using a sterile swab, the bacterial suspensions were then evenly inoculated on the surface of the Müller-Hinton agar plates (Oxoid, Basingstoke, UK). The following antibiotics, of both veterinary and human health importance, were used: amoxicillin-clavulanic acid (30μg), ampicillin (10μg), tetracycline (30μg), chloramphenicol (30μg), cefotaxime (30μg), ciprofloxacin (5μg), nalidixic acid (30μg), colistin sulphate (10μg), imipenem (10μg) and trimethoprim-sulphamethoxazole (30μg). The choice of these antibiotics was based on a list of essential drugs recommended and prioritized by WHO/OIE (19). The plates were incubated for 18-24 hours at 37°C. The zones of inhibition were read using a digital Vernier calliper and interpreted as Susceptible (S), Intermediate (I) and Resistant (R) based on the Clinical Laboratory Standards Institute recommendations (20).

### Data processing and analysis

The recorded zones of inhibition for AST were entered and analysed using WHONET software. Frequency distribution was reported for all categories as well as proportions and profiles of antibiotic resistance.

## Results

### Isolation and identification of bacteria

*Salmonella sp*. and *E. coli* were the main bacteria isolates of interest. Overall, out of the 470 samples collected, 280 (59.6%) *E. coli* and four (0.9%) *Salmonella sp*. were isolated. The occurrence of the two pathogens per sample types and areas of sampling are shown in table 1 below. Out of the 212 litter samples collected from the poultry houses in the districts under study, 58.0% (123/212) *E. coli* were isolated and no *Salmonella sp*. was found. The *E. coli* was mostly isolated in Kabwe (84.6%, 33/39) and Choma (76.4%, 13/17), whilst its occurrence was low in Chongwe (39.1%, 9/23). The occurrence of *E. coli* in other districts was almost the same (Table 1).

**Table 1.** Distribution of *Salmonella spp*. and *E. coli* isolates by location

| Sampling areas | % *Salmonella* sp. (n) |  |  |  | % *E. coli* isolates (n) |  |  |  |
| --- | --- | --- | --- | --- | --- | --- | --- | --- |
|  | Litter swabs % (n) | Cloaca swabs % (n) | Carcass swabs % (n) | Total % (n) | Litter swabs % (n) | Cloaca swabs % (n) | Carcass swabs % (n) | Total % (n) |
| Abattoir A | - | 0.0 (0/30) | 0.0 (0/30) | 0.0 (0/60) | - | 66.7 (20/30) | 56.7 (17/30) | 61.7 (37/60) |
| Abattoir B | - | 4.4 (2/45) | 4.4 (2/45) | 4.4 (4/90) | - | 91.1 (41/45) | 88.9 (40/45) | 90.0 (81/90) |
| Lusaka | 0.0 (0/24) | - | - | 0.0 (0/24) | 50.0 (12/24) | - | - | 50.0 (12/24) |
| Choma | 0.0 (0/17) | 0.0 (0/35) | - | 0.0 (0/52) | 76.4 (13/17) | 31.4 (11/35) | - | 46.2 (24/52) |
| Kabwe | 0.0 (0/39) | 0.0 (0/40) | - | 0.0 (0/79) | 84.6 (33/39) | 35.0 (14/40) | - | 59.5 (47/79) |
| Kitwe | 0.0 (0/45) | 0.0 (0/33) | - | 0.0 (0/78) | 53.3 (24/45) | 42.4 (14/33) | - | 48.7 (38/78) |
| Chilanga | 0.0 (0/31) | - | - | 0.0 (0/31) | 45.2 (14/31) | - | - | 45.2 (14/31) |
| Kafue | 0.0 (0/33) | - | - | 0.0 (0/33) | 54.5 (18/33) | - | - | 54.5 (18/33) |
| Chongwe | 0.0 (0/23) | - | - | 0.0 (0/23) | 39.1 (9/23) | - | - | 39.1 (9/23) |
| Total | 0.0 (0/212) | 1.1 (2/183) | 2.7 (2/75) | 0.9 (4/470) | 58.0 (123/212) | 54.6 (100/183) | 76 (100/183) | 59.6 (280/470) |

At the abattoir, from the 150 samples collected, *E. coli* and *Salmonella sp*. were isolated at a proportion of 78.7% (118/150) and 2.67% (4/150), respectively. Out of the total *E. coli* isolates, 31.4% (37/118) were isolated from abattoir A while 68.6% (81/118) were isolated from abattoir B. All the *Salmonella sp*. isolates originated from abattoir B of which two were from cloacal swabs and two from carcass swabs.

At the open markets, out of the 108 cloacal swabs samples collected from the three districts under study, only *E. coli* were isolated (36.1%, 39/108). When considering the occurrence of *E. coli* concerning the number of samples collected from each district, the pathogen was mostly isolated in Kitwe (42.4%, 14/33).

### Antibiotic susceptibility testing

One out of the four *Salmonella sp*. isolates exhibited resistance to 3 antibiotics namely, Amoxicillin-clavulanic acid (25%, 95% CI: 1.3% - 78.1%), Ampicillin (25%, 95% CI: 1.3% - 78.1%), and Cefotaxime (25%, 95% CI: 1.3% - 78.1%). All the other isolates were susceptible to all the other antibiotics tested.

The antibiogram pattern for the 280 *E. coli* isolates revealed high sensitivity to imipenem (97.1%) and colistin sulphate (95.4%). The highest resistance was observed against tetracycline (81.4%, 95% CI: 76.2 – 85.7%) (Table 2). In all the districts where cloacal swabs were collected from open markets, isolates showed high resistance to tetracycline, with those from Choma showing the highest resistance (81.8%), followed by those from Kitwe (85.7%) and Kabwe (71.4%).

**Table 2:**
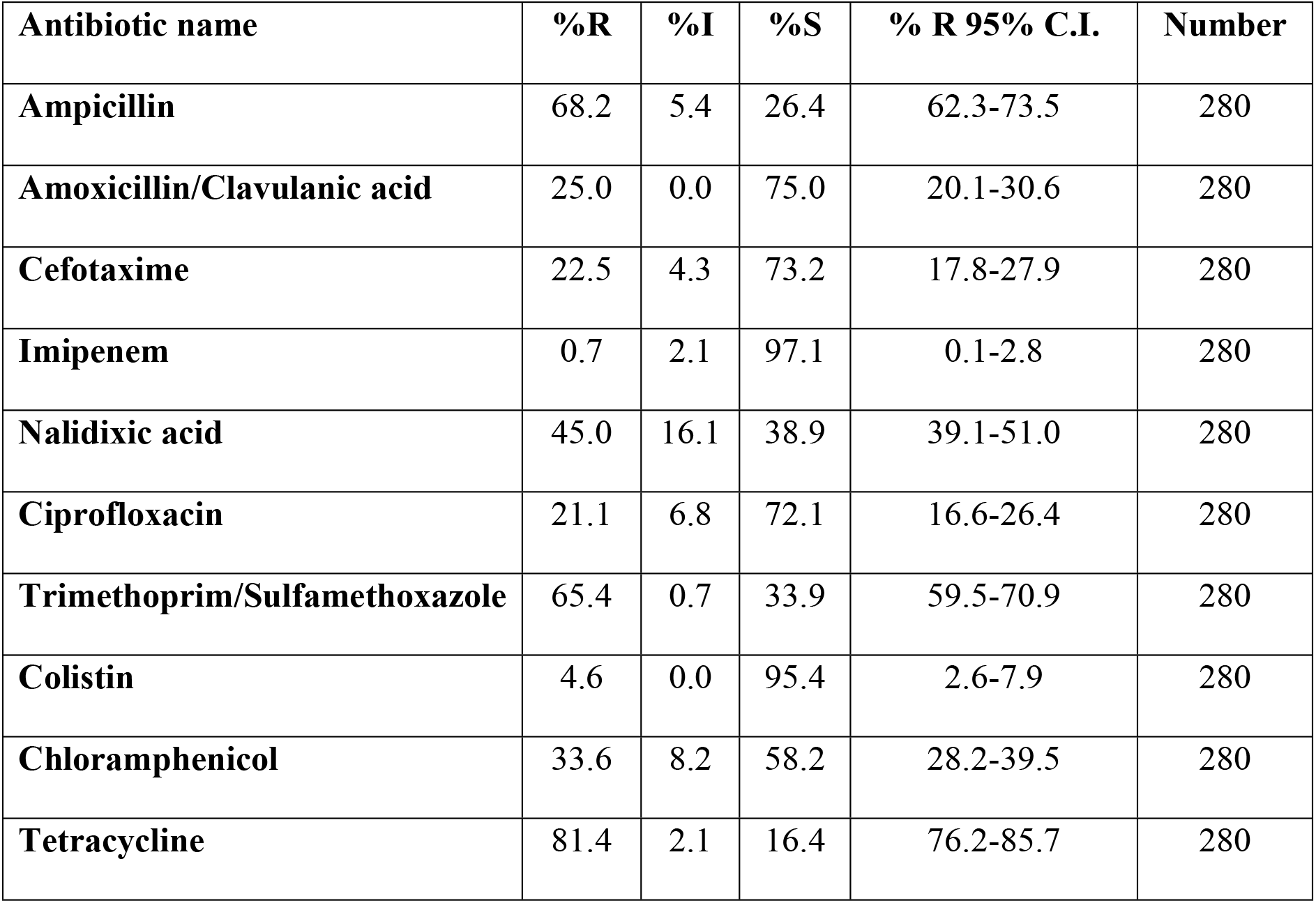
Overall antibiotic resistance patterns for *E. coli* isolates.

Isolates from the litter (poultry houses) showed very high resistance to tetracycline (91.9%, 95% CI: 85.2 – 95.8%), while least resistant was observed with imipenem (1.6%, 95% CI: 0.3 – 6.3%) (Table 3).

**Table 3:** Antibiotic resistance patterns for *E. coli* isolated from litter in all districts sampled.

| Antibiotic name | %R | %I | %S | % R 95%C.I. | Number |
| --- | --- | --- | --- | --- | --- |
| Ampicillin | 69.1 | 4.9 | 26.0 | 60.0-76.9 | 123 |
| Amoxicillin/Clavulanic acid | 36.6 | 0.0 | 63.4 | 28.2-45.8 | 123 |
| Cefotaxime | 22.0 | 5.7 | 72.4 | 15.2-30.5 | 123 |
| Imipenem | 1.6 | 2.4 | 95.9 | 0.3-6.3 | 123 |
| Nalidixic acid | 39.8 | 18.7 | 41.5 | 31.2-49.0 | 123 |
| Ciprofloxacin | 17.1 | 7.3 | 75.6 | 11.1-25.2 | 123 |
| Trimethoprim/Sulfamethoxazole | 71.5 | 1.6 | 26.8 | 62.5-79.1 | 123 |
| Colistin | 8.9 | 0.0 | 91.1 | 4.7-15.7 | 123 |
| Chloramphenicol | 34.1 | 8.9 | 56.9 | 25.9-43.3 | 123 |
| Tetracycline | 91.9 | 0.8 | 7.3 | 85.2-95.8 | 123 |

The antibiogram pattern for *E. coli* isolated from abattoirs revealed that all the 118 isolates were susceptible to colistin sulphate and imipenem (Table 4) but displayed variable resistance patterns against the other antibiotics. The majority of isolates showed high resistance against ampicillin (72.9%, 95% CI: 63.8 – 80.5%), and the least resistance against Amoxicillin/Clavulanic acid (10.2%, 95% CI: 5.6 – 17.5%).

**Table 4:**
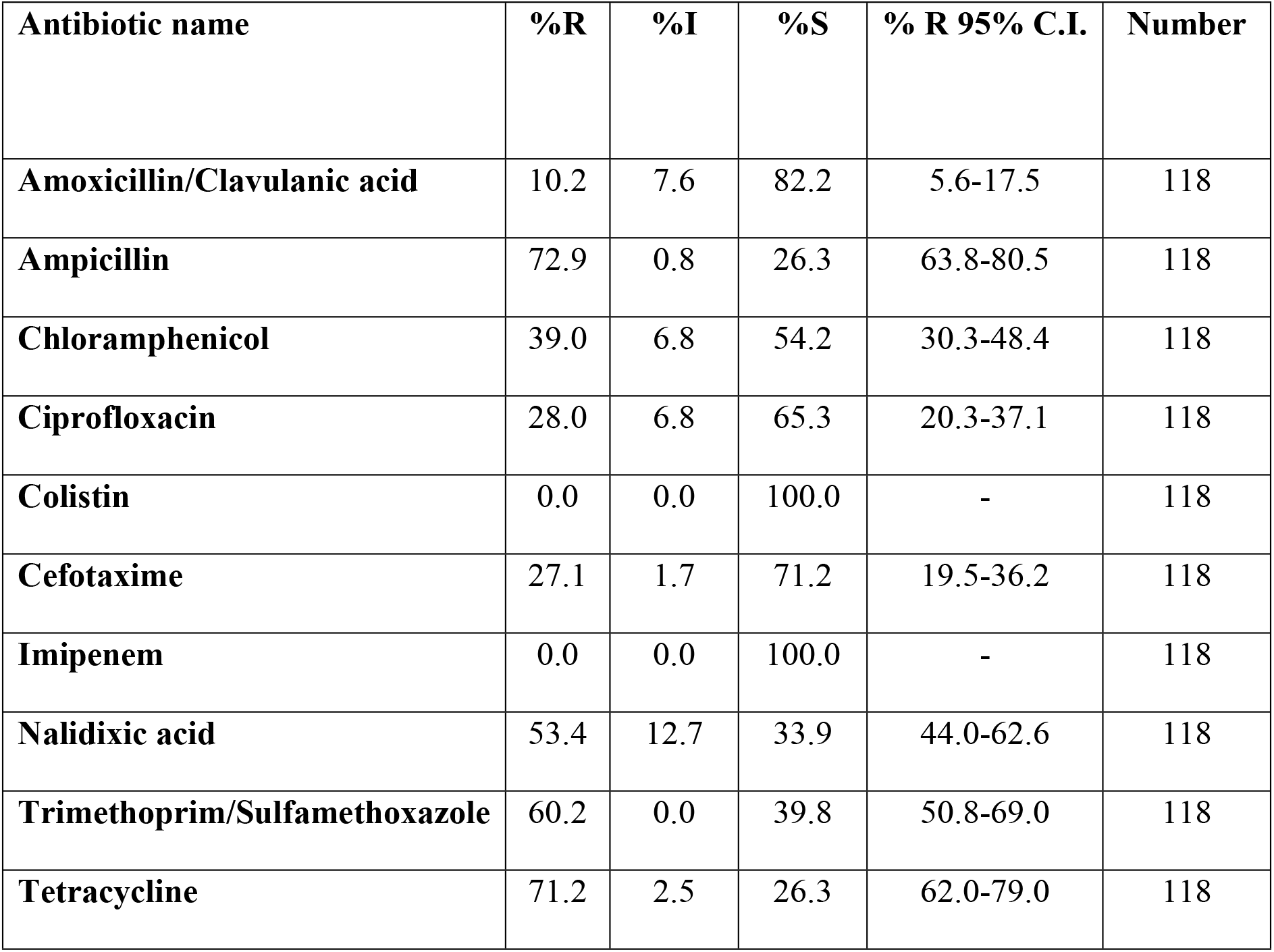
Antibiotic resistance patterns for *E. coli* isolated from cloacal and carcass swabs in abattoirs (Lusaka province).

Isolates from samples obtained from open markets in Choma, Kabwe and Kitwe showed a similar resistance pattern, with the highest resistance against tetracycline (79.5%), but no resistance to imipenem and amoxicillin/clavulanic acid (Table 5).

**Table 5:** *E. coli* Antibiogram resistance patterns of cloacal swabs samples from open markets from Choma, Kitwe and Kabwe.

| Antibiotic name | %R | %I | %S | % R 95% C.I. | Number |
| --- | --- | --- | --- | --- | --- |
| Ampicillin | 51.3 | 20.5 | 28.2 | 35.0-67.3 | 39 |
| Amoxicillin/Clavulanic acid | 0.0 | 10.3 | 89.7 | 0.0-11.2 | 39 |
| Cefotaxime | 10.3 | 7.7 | 82.1 | 3.4-25.2 | 39 |
| **Imipenem** | 0.0 | 7.7 | 92.3 | 0.0-11.2 | 39 |
| **Nalidixic acid** | 35.9 | 17.9 | 46.2 | 21.7-52.8 | 39 |
| **Ciprofloxacin** | 12.8 | 5.1 | 82.1 | 4.8-28.2 | 39 |
| **Trimethoprim/Sulfamethoxazole** | 61.5 | 0 | 38.5 | 44.6-76.2 | 39 |
| **Colistin** | 5.1 | 0 | 94.9 | 0.9-18.6 | 39 |
| **Chloramphenicol** | 15.4 | 10.3 | 74.4 | 6.4-31.2 | 39 |
| **Tetracycline** | 79.5 | 5.1 | 15.4 | 63.1-90.1 | 39 |

### Multidrug resistance and resistance profiles

Out of the 280 isolates that were subjected to susceptibility testing, 92.9% (260/280) were resistant to one or more antibiotics. Furthermore, 55% (154/280) of the *E. coli* isolates showed resistance to two or more classes of antibiotics, indicating multi-drug resistance (MDR). MDR was defined as resistance to at least one agent in at least three antimicrobial classes tested. The highest MDR profile, 20.7% (58/280) of the isolates, was to the ampicillin-tetracycline-trimethoprim/sulfamethoxazole drug combination. A small proportion, 4.3% (12/280 of the isolates) showed Extensive Drug resistance (XDR) (not susceptible to at least one agent in all but two or fewer antimicrobial categories).

## Discussion

This study found antibiotic-resistant *Salmonella sp*. and *E. coli* in broiler chickens at farm level, the abattoirs and open markets in selected districts of Zambia, which are of public health importance. No *Salmonella sp*. were isolated from either chicken litter or live chicken cloacal swabs at open markets but four *Salmonella spp*. were isolated from chickens at an abattoir in Lusaka. The *Salmonella sp*. isolated from the abattoirs in this study corroborates the findings by (21) and (22) who reported proportions of 2.6% and 2.0%, respectively. Both of the above studies were done from abattoirs in Lusaka with similar setups. However, the frequency of isolation was lower than in two previous studies conducted in Zambia, in which one reported a proportion 28% (23) and the second one reported a proportion of 16.2% (24). However, Hang’ombe et al only used biochemical tests for definitive diagnosis of *Salmonella sp*., whilst in this study, both biochemical and molecular tests were used, thereby improving the validity of the current findings.

Other studies done outside Zambia reported a high prevalence of *Salmonella sp*. at the rates of 43.6% (25) and 60.0% (26). Variation in the frequencies of isolation could be attributed to the sampling methods used in these studies, where (25) collected samples over a long period, whilst (26), sampled only at critical control points. It is reported that isolation the frequency of *Salmonella sp*. in an infected host is affected by the biological nature of the pathogen and its shedding pattern, which is seasonal and depends on environmental factors (27, 28).

This study found a high proportion of *E. coli* at abattoir level and low proportion from open markets and farms. The high isolation frequency of *E. coli* was expected since it is a commensal bacterium. However, the isolation rates from open markets and farms were lower than rates those reported in a study conducted in Spain (29). Many factors could have contributed to this, among them antibiotic usage and seasonal variation (28). The widespread antibiotic usage is the main risk factor for an increase in the occurrence of bacterial resistant strains (30). Furthermore, some studies have reported that seasonal variation affects the rate of bacterial shedding, being more in seasons of higher temperatures and less in seasons of cooler temperatures (31, 32).

The high isolation rate of *E. coli* in this study, specifically at abattoirs, corroborates with previous findings in Zambia (10, 23). These authors reported a high prevalence of *E. coli* including the presence of extended-spectrum β-lactamase (ESBL) producers. A study done in Turkey by (26) in broilers chickens destined for slaughter also found chickens to be highly contaminated with bacteria, especially with potential human pathogenic bacteria such as coliforms and *Salmonella sp*. (33). In this particular study, high contamination levels of *E. coli* on chicken carcasses were associated with carcass contamination with gut products, which occur during the process of evisceration.

*Salmonella sp*. isolates in this study showed reasonable resistance to amoxicillin-clavulanic acid, ampicillin and cefotaxime. This is closely related to what other authors have reported (15, 34, 35). Further, resistance to third-generation cephalosporins such as cefotaxime has not been often reported. The frequency and extent of *Salmonella sp*. resistance to antimicrobials drugs vary based on their usage in animal production and humans. as well as on ecological differences in the epidemiology of *Salmonella sp*. infections (36).

Low antibiotic resistance pattern of *E. coli* to imipenem and colistin sulfate was reported in this study. This could be attributed to the fact that imipenem and colistin sulfate are the last line of antibiotics for treating human bacterial infections and are not often used in food production. However, during fieldwork, the research team noticed that some farmers administered veterinary products containing colistin as an active compound. This may lead the bacteria to become resistant to this class of antibiotics due to increased exposure. On the other hand, maximum resistance was observed against tetracycline in both farm and open market samples. This was consistent with previous findings by (10), who further observed that 45.5% of the *E. coli* isolates exhibited MDR to six or more drugs. This was consistent with findings in this study in which it was observed that more than 50% of the *E. coli* isolates showed an MDR pattern, with the highest resistance profile being associated with ampicillin, tetracycline and trimethoprim/sulfamethoxazole. In other studies, MDR *Enterobacteriaceae*, including both *Salmonella spp*. and *E. coli*, have been isolated and this has been attributed to the use of growth promoters (1, 37). These findings were also consistent with those in previous study conducted in Zambia, in which it was also observed that *E. coli* isolates from cattle had high resistance against sulfamethoxazole/trimethoprim, ciprofloxacin, ampicillin and tetracycline (13).

Tetracycline has also been used as a growth enhancer and a therapeutic agent in food production (38), hence the high level of resistance observed in this study is not surprising. In Zambia, tetracycline has been used extensively to treat diseases and has given rise to the resistance of bacteria (39). Some of the major factors leading to AMR in *E. coli* include antibiotic use, overcrowding and poor sanitation (8, 40). These factors are typical of intensive poultry farming and explain the prevalence and degree of resistance in *E. coli* of poultry litter at the farms (7).

In another study, (11) found that the use of antimicrobials in veterinary practice as therapeutic and prophylactic agents, in addition to use as antimicrobial growth promoters, greatly influences the prevalence of resistance in animal bacteria and poses risk for the emergence of antibiotic resistance in human pathogens. The author further observed that isolates which are resistant to two or more antibiotics may have originated from high-risk sources of contamination like commercial poultry farms, where antibiotics are commonly used (11).

In this study, it was observed that a significant number of isolates were resistant to more than one antibiotic. This is consistent with the study by (11), which provided direct evidence that antimicrobial use in animals selects for antimicrobial-resistant bacteria that may be transferred to humans through food or direct contact with animals. This was also in consonance with previous findings in a study conducted at the University Teaching Hospital in Lusaka, Zambia, on stool samples obtained from children under the age of 5 years, in which *Salmonella sp*. and *E. coli* were also found to be multidrug-resistant (41).

### Conclusion and recommendation

This study revealed that both *Salmonella sp*. and *E. coli* are resistant to several antibiotics of both animal and human importance with similar patterns at all the three levels: farm, abattoir and open markets.

The resistance patterns in both species found in food meant for human consumption constitute a major public health concern. This study has further shown that MDR of *Salmonella sp*. and *E. coli* in broiler chickens may largely contribute to the wider and broad challenge of antimicrobial resistance and at the same time provide useful information for monitoring and surveillance purposes. The overall implication is that these resistant bacteria may be transmitted to humans and may end up causing treatment failures leading to increased morbidity and mortality. More studies need to be done on the abattoir workers (hands and faecal samples) to gain insight into their possible contribution to poultry meat AMR bacteria contamination.

## Acknowledgements

We would like to thank the World Bank-funded African Centre of Excellence in Infectious Diseases of Humans and Animals (ACEIDHA) Project and the CAPAZOMANITECO Project at the University of Zambia for the support provided during this study. We would like to thank Mr. Joseph Ndebe, Mr. Penjani Kapila, Dr. Alfred Mangani, Ms. Musonda Mubanga and Dr. Wiza Mwasinga for their technical support and guidance rendered during this study.

